# RecoverY: K-mer based read classification for Y-chromosome specific sequencing and assembly

**DOI:** 10.1101/148114

**Authors:** Samarth Rangavittal, Robert S. Harris, Monika Cechova, Marta Tomaszkiewicz, Rayan Chikhi, Kateryna D. Makova, Paul Medvedev

**Affiliations:** Department of Biology, Pennsylvania State University, University Park, Pennsylvania 16802, USA; CNRS, CRIStAL, 59655 Villeneuve d’Ascq, France.; Department of Computer Science and Engineering, Pennsylvania State University, University Park, Pennsylvania 16802, USA.; The Center for Computational Biology and Bioinformatics, Pennsylvania State University, University Park, Pennsylvania 16802, USA.; Department of Biochemistry and Molecular Biology, Pennsylvania State University, University Park, Pennsylvania 16802, USA.

## Abstract

**Motivation:** The haploid mammalian Y chromosome is usually under-represented in genome assemblies due to high repeat content and low depth due to its haploid nature. One strategy to ameliorate the low coverage of Y sequences is to experimentally enrich Y-specific material before assembly. Since the enrichment process is imperfect, algorithms are needed to identify putative Y-specific reads prior to downstream assembly. A strategy that uses *k*-mer abundances to identify such reads was used to assemble the gorilla Y (Tomaszkiewicz et al 2016). However, the strategy required the manual setting of key parameters, a time-consuming process leading to sub-optimal assemblies.

**Results:** We develop a method, RecoverY, that selects Y-specific reads by automatically choosing the abundance level at which a k-mer is deemed to originate from the Y. This algorithm uses prior knowledge about the Y chromosome of a related species or known Y transcript sequences. We evaluate RecoverY on both simulated and real data, for human and gorilla, and investigate its robustness to important parameters. We show that RecoverY leads to a vastly superior assembly compared to alternate strategies of filtering the reads or contigs. Compared to the preliminary strategy used in Tomaszkiewicz et al (2016), we achieve a 33% improvement in assembly size and a 20% improvement in the NG50, demonstrating the power of automatic parameter selection.

**Availability:** Our tool RecoverY is freely available at https://github.com/makovalab-psu/RecoverY

**Supplementary information:** Attached as an additional file.

## 1 Introduction

The haploid mammalian Y chromosome reference sequence is often not properly assembled as part of large next generation sequencing (NGS) projects, for several reasons. The Y is absent from the female and only present in one copy in the male. Therefore, to obtain the desired sequencing depth, twice as much sequencing needs to be performed. Further, the presence of repeat families shared with the autosomes and the presence of regions with high homology to the X chromosome complicate the identification and assembly of the non-unique regions of the Y (reviewed in Tomaszkiewicz et al 2017).

There are several targeted approaches for assembling the Y. Single-haplotype iterative mapping and sequencing (SHIMS) is a BAC-based technique which was used to generate assemblies of the human (Skaletsky et al 2003), chimpanzee (Hughes et al 2010), rhesus macaque (Hughes et al 2012), and mouse Y chromosomes (Soh et al 2014). While SHIMS remains a highly accurate technique, it is cost- and time-prohibitive for most projects. Alternatively, deep NGS sequencing of a male can produce an assembly that includes Y chromosome sequence. The challenge in this case is to identify which of the contigs originate from the Y. Contigs that do not align to the female assembly of the same species (if available) can be flagged as coming from the Y. Alternatively, the reads can be filtered using such alignments prior to assembly. However, such approaches are deficient at identifying Y sequence that has homology to the X, such as the pseudo-autosomal regions, or to the autosomes, such as the *DAZ* gene region (Saxena et al 2000).

Because of the different copy count in males (XY) vs females (XX), male-specific sequences can also be identified based on the number of reads aligning to them (i.e. their sequencing depth). Sequencing both a male and female of a species and comparing the read depth along the male assembly can help identify Y contigs. Methods based on this approach include the Y chromosome genome scan (Carvalho and Clark 2013) and the chromosome quotient method (Hall et al 2013). Such approaches still require sequencing the male at high-depth and additional sequencing of the female. Moreover, they may still mis-classify high-copy transposable elements and other high-copy number regions shared with the X or autosomes.

In order to decrease the cost of sequencing, experimental techniques have been developed to increase the amount of Y chromosome material present in the sample. This process of chromosome-specific enrichment can be achieved by the experimental techniques of flow sorting (Dolezel et al 2012) or microdissection (Zhou and Hu, 2007). In particular, flow sorting is a separation process by which a chromosome of interest can be separated using its size and AT/GC ratio (Dolezel et al 2012). Flow sorting has been applied prior to sequencing and assembly of the Gorilla Y (Tomaszkiewicz et al 2016) and the pig sex chromosomes (Skinner et al 2016).

Although flow sorting increases the amount of Y chromosome data, there still remains a significant amount of non-Y sequence in the sample. This may originate from debris from large chromosomes, or from chromosomes that have similar size and GC content to the Y (e.g. chromosomes 21 and 22 in the human and great apes). Consequently, the read data will consist of genome-wide reads with an enrichment for Y reads at a level dictated by the efficiency of flow sorting. This non-specificity must then be removed from the data, either before or after assembly. This can be accomplished by aligning the reads or the contigs against a known female assembly, as described above for deep NGS male sequencing. However, as already mentioned, such strategies require a female reference and are inefficient in retaining regions with high homology to the X chromosome and autosomes.

An alternative strategy of isolating Y-specific reads was originally proposed by our group in Tomaszkiewicz et al. (2016). This is an alignment-free strategy that uses *k*-mer (substring of length *k*) coverage as a discriminator to identify reads originating from the Y. The underlying principle is that, in the case of an enriched data set, *k*-mers originating from the Y chromosome occur at a higher abundance than non-repetitive *k*-mers from elsewhere in the genome. *K*-mers that have an abundance above a user-selected threshold are selected as *Y-mers* (Y-specific k-mers). Subsequently, reads that contain a number of Y-mers above a user-defined threshold (i.e. the *Y*-mer match threshold) are identified as originating from the Y and used to perform an assembly. This strategy was used to assemble the gorilla Y chromosome from flow-sorted data, however, it required manually finding both the abundance and *Y*-mer match thresholds using time-consuming and potentially biased guess-and-check approaches.

In this paper, we present an improved and automated tool to isolate Y chromosome specific reads (RecoverY). Its main improvement over the original strategy proposed in Tomaszkiewicz et al. (2016) is an algorithm to automate the choice of the abundance threshold and an automated calculation of the Y-mer match threshold. These are executed prior to the filtering step of RecoverY. We evaluate the accuracy of these thresholds and the effect of the choice of *k* using simulated data from the human Y. Finally, we run RecoverY on the simulated data and on the real flow-sorted Gorilla Y data. We demonstrate that the resulting assemblies are superior to the ones obtained with the alternate approaches of using alignment to the female to either pre-filter the reads or post-filter the contigs. These improvements also lead to a longer and more contiguous assembly of the gorilla Y than using the parameters proposed in Tomaszkiewicz et al (2016). RecoverY is open-source and freely available on https://github.com/makovalab-psu/RecoverY.

## 2 Methods

Figure 1 shows the workflow for sequencing and assembling the Y chromosome using RecoverY. First, the DNA is enriched for the Y chromosome by flow-sorting. Then, it is sequenced using Illumina technology, generating what we call fsY reads. Then, RecoverY is run to identify Y-specific reads. Finally, these reads are passed downstream to the assembler. In the following, we describe the RecoverY method.

**Fig. 1.**
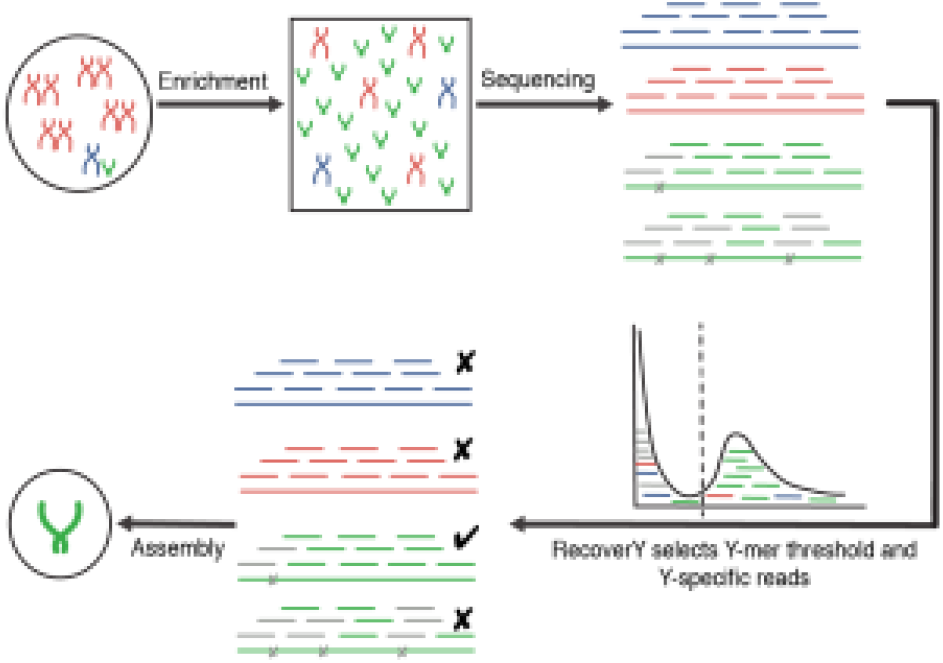
Workflow for sequencing and assembly of the Y chromosome with RecoverY. Y-enrichment increases the proportion of chromosome Y (in green) as compared to autosomes (in red) or chromosome X (in blue). Subsequent sequencing results in an increased coverage of Y. Four reads are shown here for illustrative purposes, along with their constituent *k*-mers. Grey cross marks within reads indicate sequencing errors, which affect (grey) *k*-mers overlapping the erroneous base. These error *k*-mers, along with non-Y *k*-mers, fall mostly to the left of the abundance threshold selected by RecoverY, shown by the vertical black dotted line in the histogram. Finally, RecoverY identifies as originating from the Y only those reads which have a high number of constituent *k*-mers with an abundance above the threshold. These reads are then used for downstream assembly.

To minimize time and memory requirements when running on large paired-end data sets, by default each step during the RecoverY process is run only on the forward (i.e. left or R1) reads in the data set. Once all the Y-specific forward reads are selected by the RecoverY procedure, the corresponding reverse (right or R2) reads for these pairs are included in the output data set.

The first step of RecoverY is to count the occurrences of different *k*-mers in the fsY reads. This can be performed using any *k*-mer counting software (e.g., BFcounter (Melsted and Pritchard, 2011), Jellyfish (Marcais and Kingsford, 2011), KMC3 (Kokot et al 2017), and khmer (Crusoe et al 2015)). RecoverY uses DSK v2.0.2 (Rizk et al 2013). As most of the very low abundance *k*-mers are erroneous, we use the minimum threshold recommended by DSK (-abundance-min 3) to immediately discard these k-mers. This significantly reduced the computational resources used by downstream steps.

Next, the user must provide a set of sequences which are expected to occur in single copy on the Y chromosome. The *k*-mers from these sequences are called ‘trusted *k*-mers.’ The counts of trusted *k*-mers in fsY data can act as a proxy for expected counts of Y-chromosome *k*-mers, thus aiding in determining the optimal abundance threshold. Trusted *k*-mers can be obtained from the sequence of single-copy Y chromosome genes (e.g. X-degenerate genes) from the same species -- often, these are known in advance using targeted approaches (Goto et. al. 2009). Another source of trusted *k-*mers is from single-copy sequences from a well-assembled Y chromosome of a closely related species.

More specifically, RecoverY first loads the result of *k*-mer counting into a dictionary called AllKmerCounts. Next, it creates a TrustedKmer list, by extracting all *k*-mers from the set of provided single-copy Y sequences. Subsequently, RecoverY looks up every *k*-mer from the TrustedKmer list in the AllKmerCounts dictionary, and assigns the corresponding abundance from the dictionary to this *k*-mer. The results are stored in a list and the 5th percentile of the abundances in this list are calculated. This abundance value is chosen as the abundance threshold.

RecoverY then generates the *Y*-mer table, which is a dictionary that contains all the *k*-mers in AllKmerCounts whose abundance is above the abundance threshold. The idea is that this table contains at least 95% of all the sequenced *k*-mers that originated from the Y. The *Y*-mer table allows an efficient check of whether a given *k*-mer is abundant. Next, for every input read in our data set, we consider its constituent *k*-mers. If a sufficient number of these constituent k-mers are present in the Y-mer table, we classify this read as a Y-read.

The minimum number of constituent *k*-mers, which we will refer to as the Y-mer match threshold (YT), depends on read length and error rate. RecoverY chooses the *Y*-mer match threshold automatically using the formula 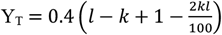. The formula is derived using the following logic. A read contains *l*−*k*+1 *k*-mers. A single error affects up to *k k*-mers -- that is, these *k*-mers are no longer in the *Y*-mer table. We aim to recover reads that have at most two errors per 100 bp window. We choose this threshold because reads with multiple errors will produce *k*-mers that, even if an error-correction algorithm is later used, are unlikely to be very informative for the construction of de Bruijn graphs in the downstream assembly process. We also tolerate that up to 60% of those *k*-mers may not be in the Y-mer table, due to undersampling. This number was chosen based on our simulation results.

## 3 Results

### 3.1 Data sets

We used both simulated and real data to evaluate RecoverY (Table 1). Two simulated data sets were generated to test different levels of enrichment. We simulated reads from the human reference genome (hg38) using the wgsim simulator v0.3.1 from the Samtools package (Li et al 2009), with default parameters for autosome sampling -- mutation rate 0.1, base error rate 0.02, fraction of indels 0.15 -- and the following parameters for X- and Y-sampling: mutation rate 0.0, base error rate 0.02, fraction of indels 0.0. The settings for mutation rate and fraction of indels reflect that the X and Y chromosomes are haploid in a male. For the first simulated data set, reads were sampled at 150x, 3x, and 6x sequencing depth from the Y, the X, and the autosomes, respectively. For the second simulated data set, reads were sampled at 300x, 3x, and 6x, respectively. These two cases simulate a male genome data set where the Y is enriched by 40.57% and 68.06%, respectively (Sup. Note 2). We refer to these data sets as human 6A_150Y and human 6A_300Y, respectively. In addition, we used a real data set of flow-sorted gorilla Y data stored under SRA accession number SRX1160374 (Tomaszkiewicz et al 2016).

**Table 1.** Simulated and real data sets used to test RecoverY

| Data set | Type | Read length | Total # reads |
| --- | --- | --- | --- |
| Human (6A_150Y) | Simulated | 150 bp | 143,272,798 |
| Human (6A_300Y) | Simulated | 150 bp | 170,831,998 |
| Gorilla fsY | Real | 150 bp | 279,209,084 |

### 3.2 K-mer abundance threshold estimation

We first applied RecoverY to select the abundance threshold for the simulated and real data sets, using a value of *k* = 25. We experimented with two lists of trusted *k*-mers. The first, called *trusted-gene-kmers*, was generated from the set of single-copy Y chromosome X-degenerate gene sequences of human, retrieved from Ensembl (Table S1). The second, called *trusted-singleton-kmers*, was based on the set of *k*-mers present in single copy in the human reference Y chromosome (hg38). To obtain the *trusted-singleton-kmers* set, these single copy *k*-mers were further filtered by removing *k*-mers that mapped to the autosomes or the X chromosome of the hg38 reference, using the BWA aligner (Li & Durbin 2009). Such *k*-mers are removed because they are too similar to non-Y *k*-mers and may negatively affect the choice of abundance threshold.

Figure 2 presents the abundance histograms generated by RecoverY, which the corresponding distributions shown in Sup. Figure 2. Note that the shape of the fsY curve closely follows that of the two trusted *k*-mer distributions except for the low-abundance *k*-mers, which is consistent with the expected effect of sequencing errors. This is the case even for the gorilla, despite the trusted *k*-mers being generated from human sequence. For the gorilla data set, RecoverY recommended a threshold of 71x when using the *trusted-gene-kmers* set and 111x using the *trusted-singleton-kmers* set. When the *trusted-gene-kmers* and *trusted-singleton-kmers* thresholds differ, we recommend choosing the lower of the two thresholds to achieve higher sensitivity. For the human, applying RecoverY resulted in a threshold of 31x for the 6A_150Y data set and 66x for the 6A_300Y data set. These thresholds were the same regardless of whether *trusted-gene-kmers* or *trusted-singleton-kmers* were used as the trusted *k*-mer set. We note that for the human dataset, using *trusted-singleton-kmers* is not a realistic experiment, since the whole human assembly was used to construct *trusted-singleton-kmers*. We include the result here only for completeness.

**Fig. 2.**
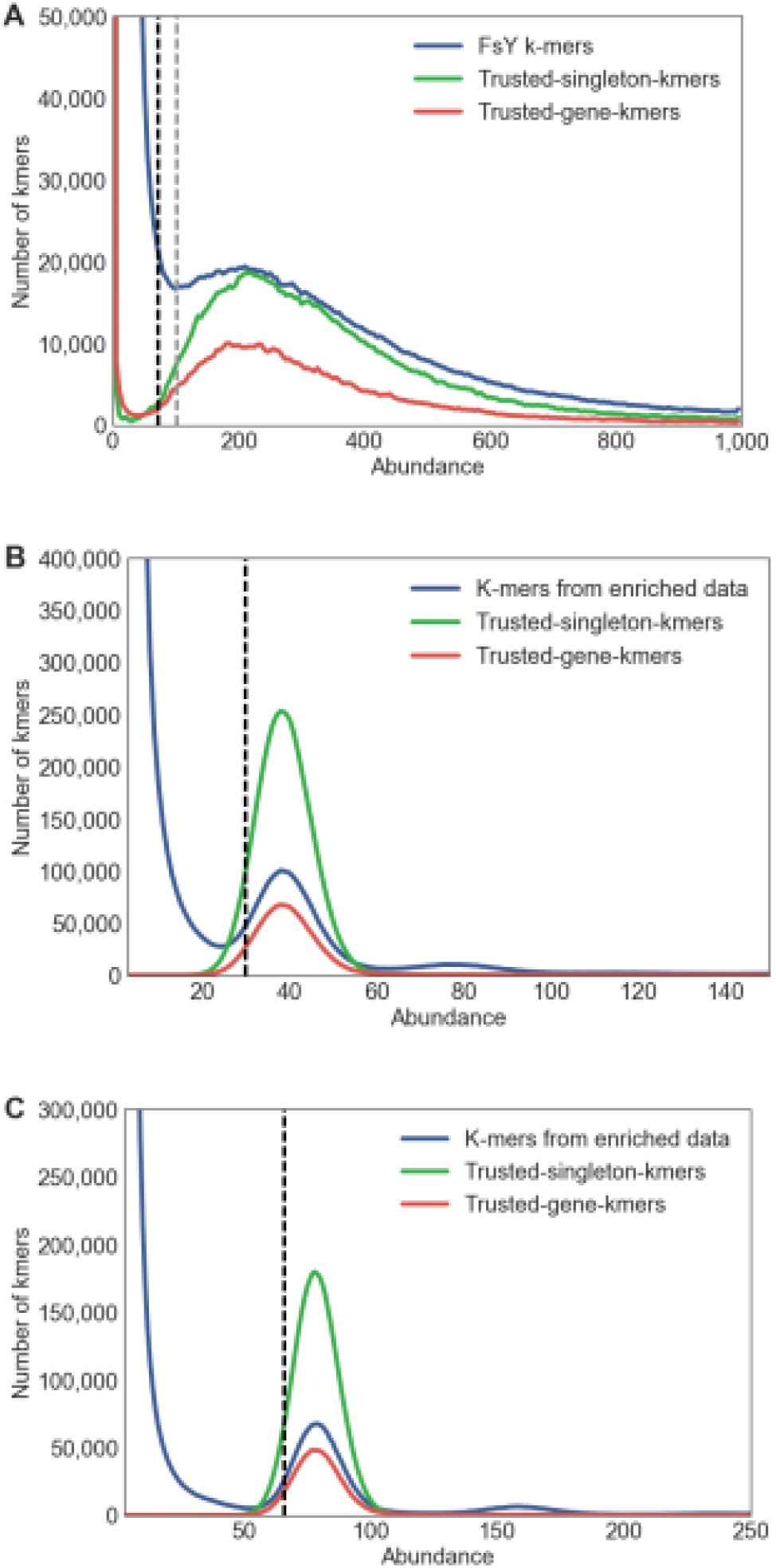
*K*-mer abundances in the gorilla flow-sorted Y real data (panel A), human male simulated data at 150x sequencing depth of the Y (6A_150Y) (panel B), and human male simulated data at 300x sequencing depth of the Y (6A_300Y) (panel C). The blue *k*-mers show the abundance histogram of *k*-mers from all the reads. We used a random subsample of 10% of the *k*-mers, effectively lowering the height of the blue curve ten-fold and making its shape visually comparable to the other curves. The green (respectively, red) curve shows the abundance histogram of *k*-mers in the dataset based on unique regions (respectively, single-copy genes). The threshold found by RecoverY using trusted-gene-kmers is shown as a dotted black vertical line. The threshold found when using trusted-singleton-kmers is the same in the case of the simulated data, but is shown as a dotted grey vertical line for the gorilla. The same data plotted as a distribution, instead of a heuristic, is shown in Sup. Figure 2.

Next, we determined the accuracy of the Y-mer table in identifying Y chromosome *k*-mers, in the two simulated data sets (Table 2). We tested the membership of all *k*-mers in the Y-mer table and in the human Y chromosome reference. Table 2 shows that RecoverY correctly identifies about 95% of all *k*-mers on the human Y chromosome reference. Of the *k*-mers from non-Y chromosomes, >99% are classified as such by RecoverY. We observe that additional coverage gives the algorithm the power to reduce false positives, without a significant effect on the false negatives.

**Table 2.** Accuracy of the Y-mer table in correctly identifying *k*-mers from the Y chromosome.

| Data set | Specificity | Sensitivity |
| --- | --- | --- |
| Human (6A_150Y) | 99.38 % | 95.32 % |
| Human (6A_300Y) | 99.79 % | 94.94% |

### 3.3 Accuracy of RecoverY in identifying Y-reads

Next, we measured the accuracy of RecoverY at classifying reads’ origin. Figure 3 shows that for every read, the number of its constituent *k*-mers that appear in the Y- mer table can be used to separate Y-origin reads (reads simulated from a Y location) from non-Y reads. However, there is some overlap of the two histograms, which naturally leads to classification errors. We measured the sensitivity as the percentage of Y-origin reads that satisfy the *Y*-mer match threshold. The specificity is the percentage of non-Y reads that do not satisfy the *Y*-mer match threshold. Figure 4 shows the tradeoff in accuracy when varying *Y*-mer match thresholds are used. Using a cutoff of 20, as suggested by our *Y*-mer match threshold formula, RecoverY achieves a sensitivity of 0.98 and a specificity of 0.83 for the 6A_300Y data set and a sensitivity of 0.98 and a specificity of 0.81 for the 6A_150Y data set. Notice that for high sensitivity values, higher coverage leads to a better specificity for the same sensitivity.

**Fig. 3.**
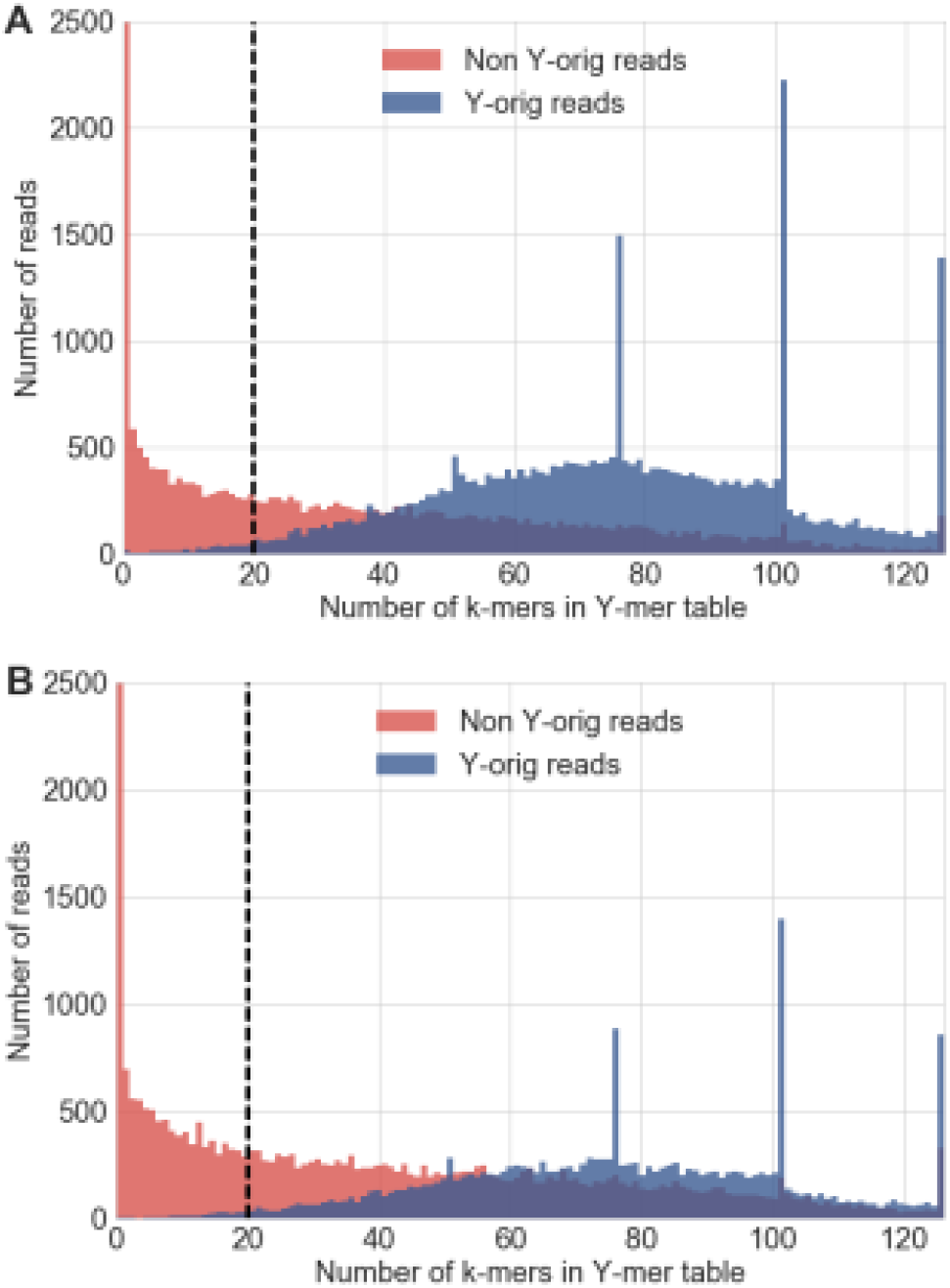
Ability of RecoverY to identify read origin. A histogram showing the numbers of reads according to their number of constituent *k*-mers in the Y-table. Panel A shows the 6A_300Y data set and panel B shows the 6A_150Y data set. The dotted black vertical line indicates the *Y*-mer match threshold chosen by RecoverY’s formula (20). Note that the first bar at *x* = 0 has been vertically cut off in the plot but has the number of reads as 49,040 for 6A_300Y and 56,935 for 6A_150Y. The spikes at *x=* 51, 76, 101 and 126 represent reads with 3, 2, 1 and 0 single base pair errors, respectively. Each plot is generated for a random subsample of 100,000 forward reads.

**Fig. 4.**
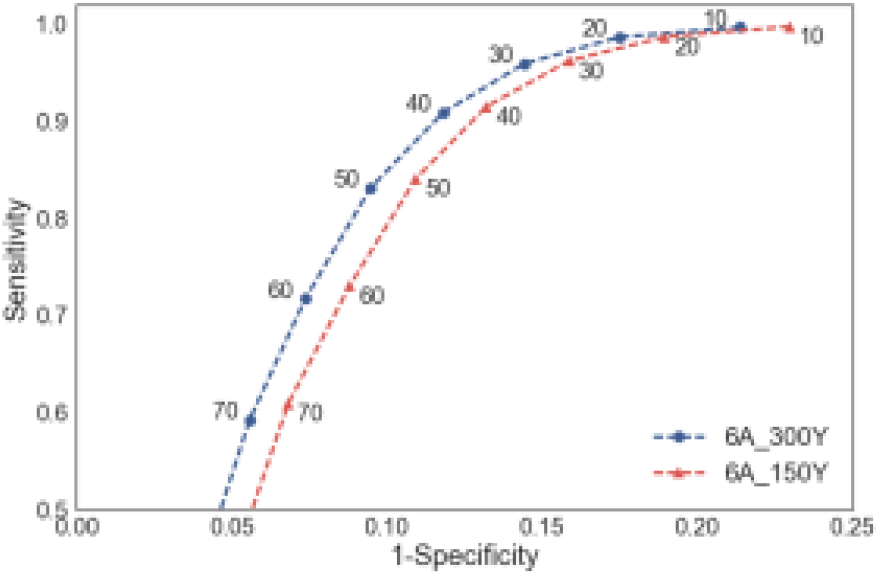
Receiver operating characteristic (ROC) plot with varying Y-mer match threshold (YT). For the same subsample of reads as used for Figure 3B, we show sensitivity and specificity for correctly identifying reads from the Y, across different YT values (intervals of 10).

We also investigated the source of false negative reads. A manual inspection (Sup. Note 1, Sup. Figure 1) suggests that the major source are reads with multiple sequencing errors. Among the constituent *k*-mers of such reads, there is a large number of very low-abundance error *k*-mers. These error *k*-mers are not in the *Y*-mer table, and such reads will be mistakenly classified as non-Y.

### 3.4 Effect of varying *k*-mer size on classification of Y-reads

In bioinformatics applications, the *k*-mer size is often a parameter that has to be estimated by trial and error. Choosing a large *k* ensures a higher proportion of unique *k*-mers in the data set at the cost of a higher probability of a *k*- mer containing an error. Conversely, choosing a too short *k* results in many repeated *k*-mers in the data set. We tested different *k*-mer lengths and constructed ROC curves for the human 6A_150Y data set (Figure 5). For each value of *k*, the abundance threshold was chosen by RecoverY and *Y* - mer match threshold was varied from 10 to 120 in increments of 10. The plot shows that with higher values of *k*, a better specificity can be achieved for a fixed sensitivity. However, higher values of *k* (*k* > 31) make it impossible to achieve high sensitivity values even at very low *Y*-mer match thresholds (Y_T_ = 10). Based on these results, the optimal choice of *k* for applying RecoverY on mammalian Y chromosomes, with similar levels of coverage and enrichment, is between 21 and 31.

**Fig. 5.**
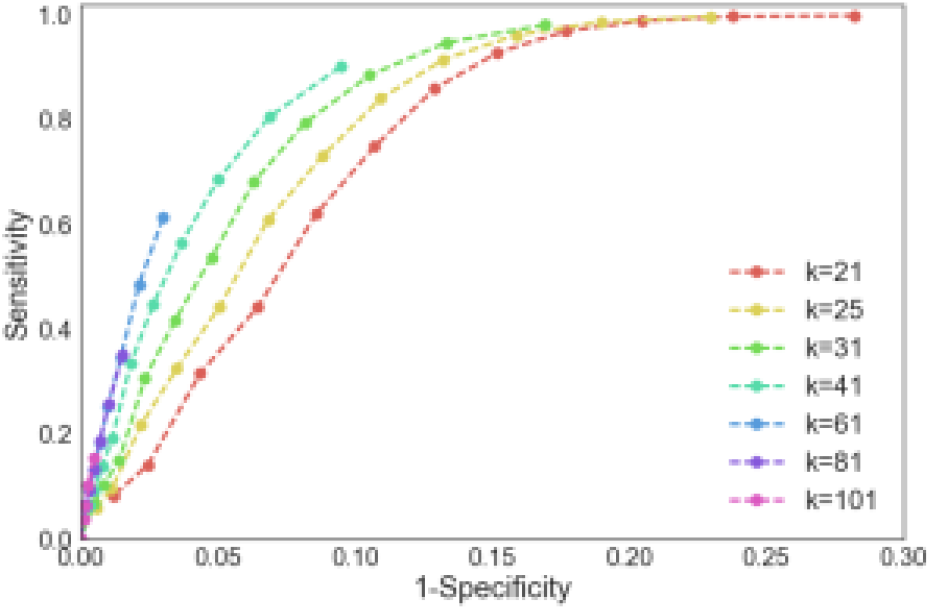
ROC plot with varying *k*-mer size. For the same subsample of 6A_150Y reads as used for Figure 3, the *k*-mer size was varied from *k* = 21 (red) to *k* = 101 (violet). The abundance thresholds as generated by RecoverY were 35 (*k*=21), 31 (*k*=25), 25 (*k*=31), 18 (*k*=41), 9 (*k*=61), 4 (*k*=81) and 4 (*k*=101). Within each colored line, solid point markers represent increasing *Y-*mer match thresholds from 10 (top right) up to a maximum threshold of 120 (bottom left), in stepwise increments of 10.

### 3.5 Effect of RecoverY on assembly

We compare the effect of RecoverY on downstream assembly quality against two alternate approaches. The first assembly, called PreFiltReads, is generated by assembling only those reads which remain unmapped to a repeat-masked female reference genome (hg38 minus the Y chromosome). RepeatMasker version open-4-0-3 was used with -s sensitive setting. BWA-MEM v0.7.5a (Li H. 2013) was used for read mapping, with default parameters (seed length 19, mismatch penalty 4, and gap open penalty of 6). The second assembly, PostFiltCtgs, is generated by assembling all reads but retaining only those contigs which remain unmapped to a repeat-masked hg38 female genome. For alignment of contigs, BLASR (Chaisson and Tesler 2012) was used with default parameters (min. seed length 12, mismatch penalty 6), and the - unaligned option to collect unaligned contigs. Among the various short read assemblers, we chose the SPAdes assembler for its ability to deal with uneven coverage profiles (Bankevich et al. 2012). We use SPAdes version 3.6.1 with the following parameters: --only-assembler and -k 21, 33, 55. Note that the *k*-mer size used in RecoverY is an independent parameter to the *k*-mer sizes used for assembly and both values might differ widely, depending on the data set. We also used Discovar assembler version r52488 (Weisenfeld et al. 2014) to replicate our conclusions on real data. Because it did not perform as well as SPAdes, we only used SPAdes for the simulated data. Soapdenovo2 (Luo et al 2012) or Minia (Chikhi et al 2013) or other assemblers could also be tried, though we did not test them on our datasets.

To evaluate the assemblies, we use the standard QUAST tool v3.1 (Gurevich et al 2013), which reports contiguity and quality metrics. It defines the NG50 (respectively, N50) as the length at which contigs of size longer or equal to that length sum to up at least half of the reference (respectively, assembly) length. It defines a misassembly as a position in a contig where flanking sequences do not align concordantly to the reference (in this case, the hg38 human Y chromosome). The number of mismatches per 100 kb is defined as the number of single-nucleotide mismatches between contigs and the reference, including SNPs and sequencing errors. Because SPAdes produces a very large number of small contigs, we filter out all contigs shorter than 1000 bp using the QUAST parameter: --min-contig 1000.

Table 3 compares the PreFiltReads and PostFiltCtgs assemblies against the one produced by assembling the RecoverY filtered reads, on the 6A_150Y dataset. Pre-filtering of reads significantly reduces the size of the data set input to the assembler, thus improving run time and memory usage. However, the assembly is much smaller than expected (~25 Mb, according to the known euchromatic length of the reference hg38 human Y assembly) (Skaletsky et al 2003). Additionally, it exhibits low N50, along with significant number of misassemblies and mismatches. Post-filtering of contigs produces an assembly that is only a tiny fraction (2%) of the expected size. Compared to these strategies, RecoverY works best. It achieves the largest assembly by far, with relatively few misassemblies and mismatches. Its N50 is five times higher than the N50 with the pre-filtering strategy, though at the expense of memory usage and speed.

**Table 3.**
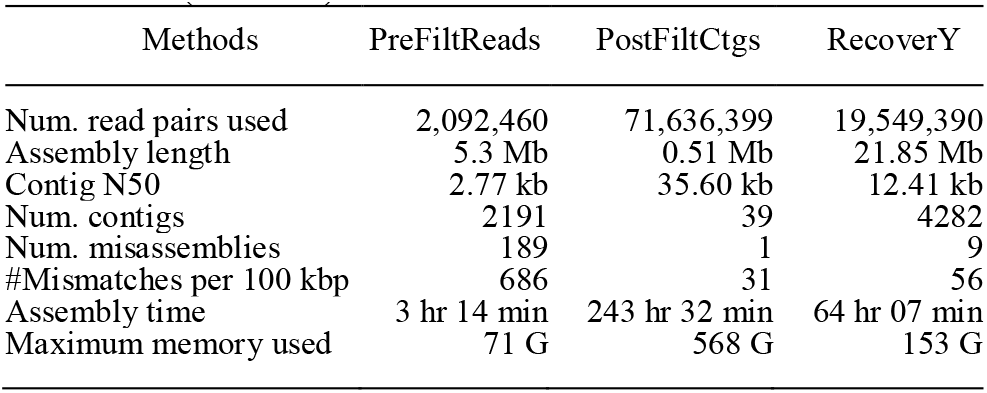
Comparison of different assembly strategies on simulated human reads (6A_150Y).

| Methods | PreFiltReads | PostFiltCtgs | RecoverY |
| --- | --- | --- | --- |
| Num. read pairs used | 2,092,460 | 71,636,399 | 19,549,390 |
| Assembly length | 5.3 Mb | 0.51 Mb | 21.85 Mb |
| Contig N50 | 2.77 kb | 35.60 kb | 12.41 kb |
| Num. contigs | 2191 | 39 | 4282 |
| Num. misassemblies | 189 | 1 | 9 |
| #Mismatches per 100 kbp | 686 | 31 | 56 |
| Assembly time | 3 hr 14 min | 243 hr 32 min | 64 hr 07 min |
| Maximum memory used | 71 G | 568 G | 153 G |

We also evaluated the improvement in assembly due to the new version of RecoverY as compared to the preliminary strategy used while assembling the draft gorilla Y chromosome (Tomaszkiewicz et al. 2016). Both versions of RecoverY were run on the gorilla fsY data set (Table 1), followed by SPAdes. For the preliminary RecoverY version, we used the parameters described in Tomaszkiewicz et al. (2016): *Y*-mer match threshold of 50 and abundance threshold of 100. For the new version, RecoverY selected an abundance threshold of 71 (Figure 2A) and a *Y*-mer match threshold of 20.

Table 4 shows that the improvements resulted in a 33% improvement in assembly size and a 20% improvement in the NG50. To evaluate the reproducibility of our findings, we repeated the experiment using the Discovar assembler version r52488 (Weisenfeld et al. 2014) in place of SPAdes (Table 5). While the absolute quality of the assembly was lower for Discovar, the relative performance of the new RecoverY to its preliminary version remained the same.

**Table 4:**
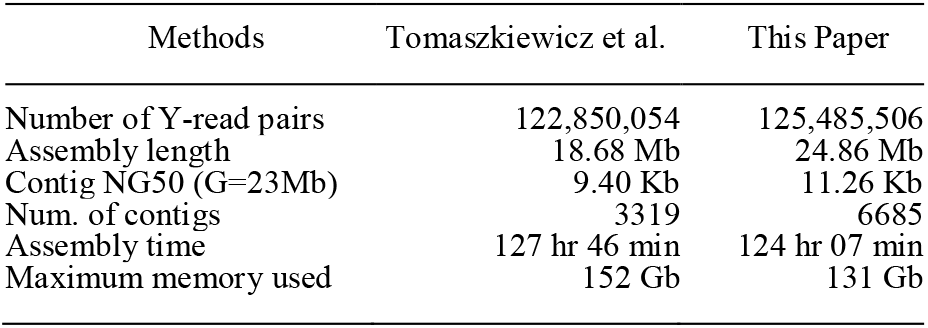
SPAdes gorilla Y assembly using the preliminary RecoverY strategy from Tomaszkiewicz et al. (2016) vs. the one proposed in this paper.

| Methods | Tomaszewicz et al. | This Paper |
| --- | --- | --- |
| Number of Y-read pairs | 122,850,054 | 125,485,506 |
| Assembly length | 18.68 Mb | 24.86 Mb |
| Contig NG50 (G=23Mb) | 9.40 Kb | 11.26 Kb |
| Num. of contigs | 3319 | 6685 |
| Assembly time | 127 hr 46 min | 124 hr 07 min |
| Maximum memory used | 152 Gb | 131 Gb |

**Table 5.** Discovar gorilla Y assembly using the preliminary RecoverY strategy from Tomaszkiewicz et al. (2016) vs. the one proposed in this paper.

| Methods | Tomaszewicz et al. | This Paper |
| --- | --- | --- |
| Number of Y-read pairs | 122,850,054 | 125,485,506 |
| Assembly length | 5.57 Mb | 8.34 Mb |
| Contig NG50 (G=8.34M) | 1489 bp | 2124 bp |
| Num. of contigs | 2655 | 3917 |
| Assembly time | 3 hr 25 min | 3 hr 45 min |
| Maximum memory used | 278.33 Gb | 281.51 Gb |

We also compare the performance of RecoverY against a non-enrichment based approach which simply sequences a male genome, assembles it, and then flags contigs that do not align to the female as being putatively from the Y (Sup. Note 3). The cumulative length of these Y-contigs is 1.51 MB with an N50 of 1895 bp (Sup. Note 3), which is an order of magnitude lower than the results from the RecoverY enrichment approach. Further, because this method relies on sequence matching and not on k-mer counts, it might fail to retrieve regions that show homology to the female autosomes or the X chromosome (e.g. pseudo-autosomal region and X-transposed regions).

### 3.6 Speed and memory performance

We measured the runtime and memory usage of DSK *k*-mer counting, RecoverY after DSK, and the downstream assembly, in all of our datasets (Table 6). We ran our experiments on a x86_64 system with up to 64 available AMD Opteron 6276 processors and 512 GB available memory. Using 8 processors, the run time of DSK and RecoverY is approximately 4 hours for each of the data sets. This is only about 3-5% of the cumulative times that include assembly. RecoverY’s memory usage, as well, is less than half of what the SPAdes assembler uses.

**Table 6.**
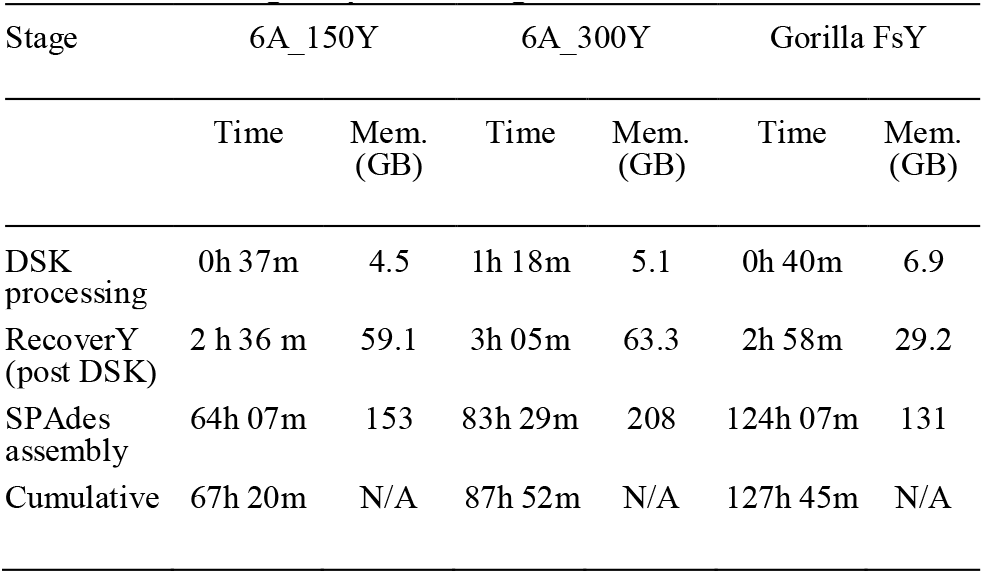
Runtime and memory usage of the different stages of RecoverY (with 8 processors). RecoverY total times are separated into DSK *k*-mer counting and post counting.

| Stage | 6A_150Y |  | 6A_300Y |  | Gorilla FsY |  |
| --- | --- | --- | --- | --- | --- | --- |
|  | Time | Mem. (GB) | Time | Mem. (GB) | Time | Mem. (GB) |
| DSK processing | 0h 37m | 4.5 | 1h 18m | 5.1 | 0h 40m | 6.9 |
| RecoverY (post DSK) | 2 h 36 m | 59.1 | 3h 05m | 63.3 | 2h 58m | 29.2 |
| SPAdes assembly | 64h 07m | 153 | 83h 29m | 208 | 124h 07m | 131 |
| Cumulative | 67h 20m | N/A | 87h 52m | N/A | 127h 45m | N/A |

A major component in the performance improvement provided by RecoverY in comparison to male whole-genome assembly based approaches is due to reduction in *k*-mer search space. For example, the raw reads that are *k*-merized in the full gorilla fsY data set result in a *k*-mer table of approximately 15 GB of space. However, after the abundance threshold is selected and the “contaminant” *k*-mers are discarded, the new Y-mer table contains only ~500 MB of *k*-mers. This represents a reduction in the *k*-mer search space by about 97%.

## 4 Discussion

In this paper, we present a method for the identification of Y-specific reads from chromosome flow-sorted data. It builds on the previous strategy of Tomaszkiewicz (2016) by automatically selecting the abundance level at which a *k*-mer is deemed to originate from the Y. The major benefit of RecoverY is that it removes non-Y reads which otherwise would confound a genome assembler. Additionally, it reduces the size of the data set provided to the assembler, thus speeding up the assembly process and reducing memory requirements. Our tests indicate that RecoverY is a drastic improvement over two other alternate filtering strategies, as well as over the preliminary version in Tomaszkiewicz (2016).

The main methodological novelty of RecoverY is the method of using trusted k-mers to automate the selection of the abundance threshold. To the best of our knowledge, this is the first application of trusted k-mers to parameter selection for de novo assembly projects. In many other tools, parameter selection is often an afterthought, resulting in low-quality results. We demonstrated the power of using trusted k-mers to choose the threshold by showing that the resulting assembly is 33% larger and has 20% higher NG50 than the one generated without the use of trusted k-mers. The notion of trusted k-mers has the potential to affect bioinformatics tools more broadly, e.g. trusted k-mers could potentially be used in de novo assembly, read error correction, metagenomic analysis, or index creation for sequencing databases. They can also potentially be used to find contaminants in an enriched dataset.

RecoverY is designed to sacrifice specificity to achieve high sensitivity, in terms of classifying read origin. The downstream cost of mistakenly classifying a non-Y read as coming from the Y is not high -- these reads tend to be scattered across the genome and contribute to very short contigs during assembly. Such short contigs are usually anyway removed at later stages. On the other hand, mistakenly classifying a Y-origin read as non-Y may have the effect of breaking an otherwise long contig during assembly. Our choice of *Y*-mer match threshold is therefore designed with this in mind, as illustrated by Figures 3 and 4. In situations where specificity is nevertheless preferred over sensitivity, the user may adapt the thresholds to better suit their needs.

An extension of RecoverY can be applied as a binary classifier on any data set to isolate an enriched chromosome of interest. Further improvements in the future include improving the specificity of the method by probabilistic weighting of *k* -mers while comparing to the *Y*-mer table. An extension of RecoverY to PacBio data is also a direction of future work. Further improvements to the run time of RecoverY are likely possible by re-implementing the codebase in a lower-level language such as C/C++.

RecoverY requires some known Y single-copy sequences to choose the abundance threshold. This information might be available prior to *de novo* assembly, but sequence-composition agnostic approaches are also possible. For instance, the abundance threshold could be selected by fitting a statistical model to the histogram, similar to the approach used by KmerGenie (Chikhi and Medvedev 2014). A threshold can also be chosen using a formula based on the enrichment levels and expected error rates. We leave these approaches to future work.

## Acknowledgements

We thank Kristoffer Sahlin for useful discussions for the optimization of the algorithm. HiSeq sequencing was performed at Pennsylvania State Genomics Core Facility, University Park, Pennsylvania.

